# HDAC6 promotes self-renewal and migration/invasion of rhabdomyosarcoma

**DOI:** 10.1101/823864

**Authors:** Thao Q. Pham, Kristin Robinson, Lin Xu, Stephen X. Skapek, Eleanor Y. Chen

**Author notes:** Equal contribution to the manuscript (Thao Q. Pham, Kristin Robinson).

## Abstract

Rhabdomyosarcoma (RMS) is a devastating pediatric sarcoma. The survival outcomes remain poor for patients with relapsed or metastatic disease. Effective targeted therapy is lacking due to our limited knowledge of the underlying cellular and molecular mechanisms leading to disease progression. In this study, we used functional assays *in vitro* and *in vivo* (zebrafish and xenograft mouse models) to demonstrate the crucial role of HDAC6, a cytoplasmic histone deacetylase, in driving RMS tumor growth, self-renewal and migration/invasion. Treatment with the HDAC6-selective inhibitor, Tubastatin A, recapitulates the HDAC6 loss-of-function phenotypes. HDAC6 regulates cytoskeletal dynamics to promote tumor cell migration and invasion. RAC1, a Rho family GTPase, is an essential mediator of HDAC6 function, and is necessary and sufficient for RMS cell migration and invasion. High expression of RAC1 correlates with poor clinical prognosis in RMS patients. Targeting the HDAC6-RAC1 axis represents a promising therapeutic option for improving survival outcomes of RMS patients.

## INTRODUCTION

Rhabdomyosarcoma (RMS) is a devastating pediatric sarcoma that displays morphological and molecular evidence for incomplete myogenic differentiation. RMS is roughly divided into two major subtypes by histologic features, embryonal (ERMS) and alveolar (ARMS). Genetically, ERMS is characterized by mutations in the receptor tyrosine kinase/RAS/PIK3CA axis found in at least 90% of cases^1^, and most ARMS cases harbor the PAX3 (or PAX7)-FOXO1 fusion transcript^2^. Despite the genetic differences, the prognosis for relapsed or metastatic disease remains poor regardless of the subtype. Effective targeted therapy is lacking due to our poor understanding of events leading to relapse or metastasis of RMS.

Several studies have shed some light on some of the genes and pathways that contribute to the metastatic potential of RMS. For example, Hepatocyte Growth Factor (HGF) and Stromal-derived Factor −1 (SDF-1) regulate metastatic behavior of cMet-positive RMS cells by directing them to the lymph nodes and the bone marrow^3^, and downregulation of the MET-CXCR4 axis decreases the migration of RMS cells in response to the SDF-1 or HGF gradients *in vitro*^4^. Deficiency of tp53 in a conserved zebrafish model of ERMS increases the metastatic potential of ERMS cells^5^. Additional targets involved in the signaling pathways such as IL-4R and PlexinA1 have also been shown to promote migratory and metastatic capacity of RMS cells^6,7^. However, the cellular and molecular mechanisms leading to invasion and metastasis of RMS cells remain to be elucidated. Specifically, the molecular players involved in altering the cytoskeletal dynamics to orchestrate RMS tumor cell invasion and metastasis remain to be identified and characterized.

Cancer stemness is mainly characterized by the self-renewal capacity of cancer cells to give rise to progeny cells that recapitulate the cancer heterogeneity^8^. The therapeutic potential of targeting cancer stem cells in solid tumors (e.g. skin squamous cell carcinoma, colorectal cancer and non-small cell lung cancer) has been demonstrated in a variety of pre-clinical animal models^9–11^. The inhibitors against several pathways involved in regulating the maintenance and survival of cancer stem cells (e.g. the Wnt, Notch and Hedgehog pathways) are in various phases of clinical trials^12,13^. However, the insight into the genes and pathways that regulate the stemness of RMS remains limited. Molecular markers such as Myf5, a marker for activated muscle satellite cells, and CD133, a transmembrane glycoprotein, have been used to enrich for tumor propagating cells in pre-clinical models of RMS^14,15^. Canonical and non-canonical Wnt pathways and TP53 have been shown to modulate the self-renewal capacity of RMS tumor-propagating cells^5,16,17^. Further investigation in preclinical models is necessary to identify additional druggable targets against the cancer stemness of RMS and determine the therapeutic benefit of targeting cancer stemness in RMS.

In our previous CRISPR screen of the Histone Deacetylases (HDACs), HDAC6 is among the selected HDACs that are essential for RMS tumor cell growth^18^. Unlike the other HDACs, HDAC6 predominantly exerts its function in the cytoplasm and has been shown to interact with substrates such as Heat Shock Protein 90 (HSP90), Extracellular signal-related Kinase 1 (ERK1), KRAS, Tubulin and Cortactin to regulate diverse cellular processes such as cell migration, adhesion and growth in several normal and neoplastic cell types^19–23^. HDAC6 has been shown to either promote or suppress cancer cell invasion and metastasis depending on the cancer cell type^24,25^. Besides the studies on the role of HDAC6 in regulating the maintenance of glioma stem cells^26,27^, there is limited knowledge on the function of HDAC6 in regulating self-renewal capacity of cancer cells.

In this study, we investigate the function of HDAC6 in various RMS features utilizing both *in vitro* and *in vivo* functional assays and demonstrate that HDAC6 is essential for RMS cell growth, migration/invasion and self-renewal. RAC1, a Rho GTPase, is a key player in mediating HDAC6 function in regulating RMS cell migration/invasion. High expression of RAC1 correlates with poor clinical survival of RMS patients. Overall, targeting the HDAC6-RAC1 axis may have significant therapeutic benefits for RMS patients.

## RESULTS

### Expression of HDAC6 in RMS

We first assessed expression of HDAC6 in primary human RMS samples by immunohistochemistry. HDAC6 expression was detected primarily in the cytoplasm at variable intensity levels in most cases of ARMS (11 of 11) and ERMS (6 of 7) (Fig. 1A-F). In contrast, five cases of pediatric muscle did not express HDAC6. A panel of ERMS (RD and SMS-CTR) and ARMS cell lines (Rh5 and Rh30) also showed cytoplasmic and perinuclear expression of HDAC6 by immunofluorescence (Fig. 1G-J). Overall, our findings showed that HDAC6 is differentially expressed in RMS cells compared to normal muscle, and that HDAC6 functions predominantly in the cytoplasm of RMS cells.

**Figure 1.**
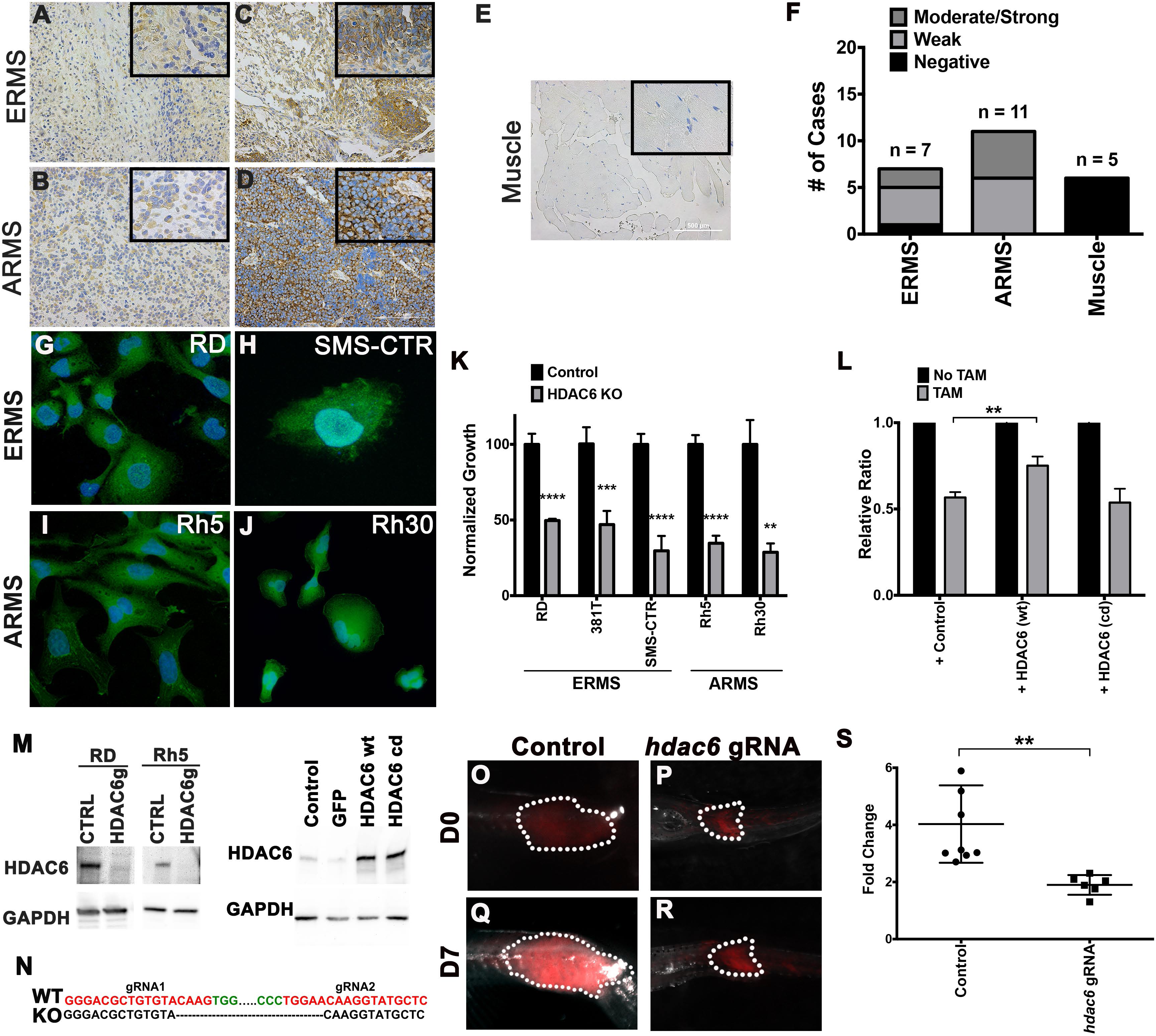
HDAC6 expression in RMS and conserved function of HDAC6 in regulating RMS tumor growth. (A-E) Immunohistochemistry for HDAC6 in representative primary ERMS and ARMS tumors and skeletal muscle tissue. Weaker staining in (A-B) and stronger staining in (C-D). No staining is seen in skeletal muscle (E). (F) Summary of IHC results in ERMS (n = 7), ARMS (n = 11) and skeletal muscle (n = 5). Immuofluorescence for HDAC6 in (G-H) ERMS cell lines (RD and SMS-CTR) and (I and J) ARMS cell lines (Rh5 and Rh30). (K) Summary of change in cell growth by cell counts following *HDAC6* knockout (KO) by CRISPR/Cas9 in a panel of RMS cell lines. The results shown are the average of 3 replicates for each cell line from one of 3 independent experiments. (L) Overexpression of GFP as control, Cas9-resistant wild-type (wt) HDAC6 and Cas9-resistant catalytically-dead (cd) HDAC6 in tamoxifen (TAM)-inducible Cas9/*HDAC6* gRNA RD line to assess change in cell growth following TAM-induction. Results shown are the average of 4 replicates from one of 3 independent experiments. (M) Western blots against HDAC6 in RD and Rh5 cells with targeted disruption of *HDAC6* by CRISPR (left panel) and in RD TAM-inducible Cas9/*HDAC6* gRNA line overexpressing GFP, wt HDAC6 and cd HDAC6 (right panel). *HDAC6*g = *HDAC6* gRNA. GAPDH was used as a loading control. (N) A snapshot of genomic DNA sequencing results demonstrating deletion in the *hdac6* locus in a CRISPR-targeted zebrafish tumor. (O-R) Representative zebrafish ERMS tumors expressing GFP control vector (O, Q) and Cas9/*hdac6* gRNA (P, R) over 7 days of growth. (S) Summary of tumor growth. n = 8 for control and n = 6 for *hdac6* gRNA-targeted ERMS tumors. Each error bar in graphs of K, L and S represents standard deviation. ** = p < 0.01; *** = p < 0.001; **** = p < 0.0001.

### Conserved role of HDAC6 in promoting tumor growth in RMS

To assess HDAC6 loss-of-function effects tumor cell growth, we generated CRISPR/Cas9-mediated *HDAC6* gene knockout in a panel of RMS cell lines (ERMS: RD, 381T, SMS-CTR; ARMS: Rh5 and Rh30). Targeted disruption of *HDAC6* resulted in a significant reduction of tumor cell growth (Fig. 1K). To assess the specificity of *HDAC6* knockout on tumor cell growth, we performed rescue experiments by overexpressing Green Fluorescent Protein (GFP) as a control, wild-type HDAC6 (modified to be Cas9-resistant) and catalytically-dead (cd) HDAC6 in RD cells harboring tamoxifen-inducible CRISPR/Cas9 targeting *HDAC6*. Overexpression of Cas9-resistant wild-type HDAC6 alleviated the *HDAC6* knockout-induced growth phenotype in comparison with overexpression of cd HDAC6 or GFP, indicating that the effect of *HDAC6* knockout on tumor cell growth is specific and that the catalytic activity of HDAC6 is required for its function in regulating RMS tumor cell growth (Fig. 1L). Depletion of HDAC6 protein from CRISPR/Cas9-mediated gene knockout and overexpression of wild-type and cd HDAC6 were confirmed by Western blots (Fig. 1M).

To assess the effects of HDAC6 loss-of-function on tumor growth *in vivo*, we generated CRISPR/Cas9-mediated *hdac6* knockout in a conserved zebrafish model of KRAS(G12D)-induced ERMS^28^ (Fig. 1N). Zebrafish ERMS tumors harboring *hdac6* knockout showed significantly reduced tumor growth by at least 50% compared to tumors harboring the GFP-scrambled gRNAs as a control (Fig. 1O-S, p < 0.01), indicating that hdac6 has a conserved role in promoting ERMS tumor growth.

### HDAC6 promotes RMS tumor growth by modulating cell cycle progression and tumor cell differentiation

To determine the tumor cell phenotypes that contributed to reduced tumor cell growth following targeted disruption of *HDAC6*, we assessed for alterations in cell cycle progression, cell death and cell differentiation in RMS cells with *HDAC6* knockout. RD and Rh5 cells with *HDAC6* knockout showed arrest in G1 or G2/M phases of cell cycle (Fig. 2A) but no significant change in programmed cell death (Fig. 2B). RMS cells are pathologically characterized by an arrest in myogenic differentiation^16,18^. We showed that targeted disruption of *HDAC6* significantly increased myogenic differentiation (approximately 2-3 folds; p < 0.01) compared to controls, based on immunofluorescence staining for myosin heavy chain (Fig. 2 C-G). RMS cells harboring targeted *HDAC6* also showed increased expression of genes (e.g. *MYOD1*, *MYH8*, *CKM*) involved in myogenesis following targeted disruption of *HDAC6* (results for RD shown in Fig. 2 H). Based on our findings, HDAC6 promotes RMS tumor growth in part by modulating cell cycle progression and tumor cell differentiation.

**Figure 2.**
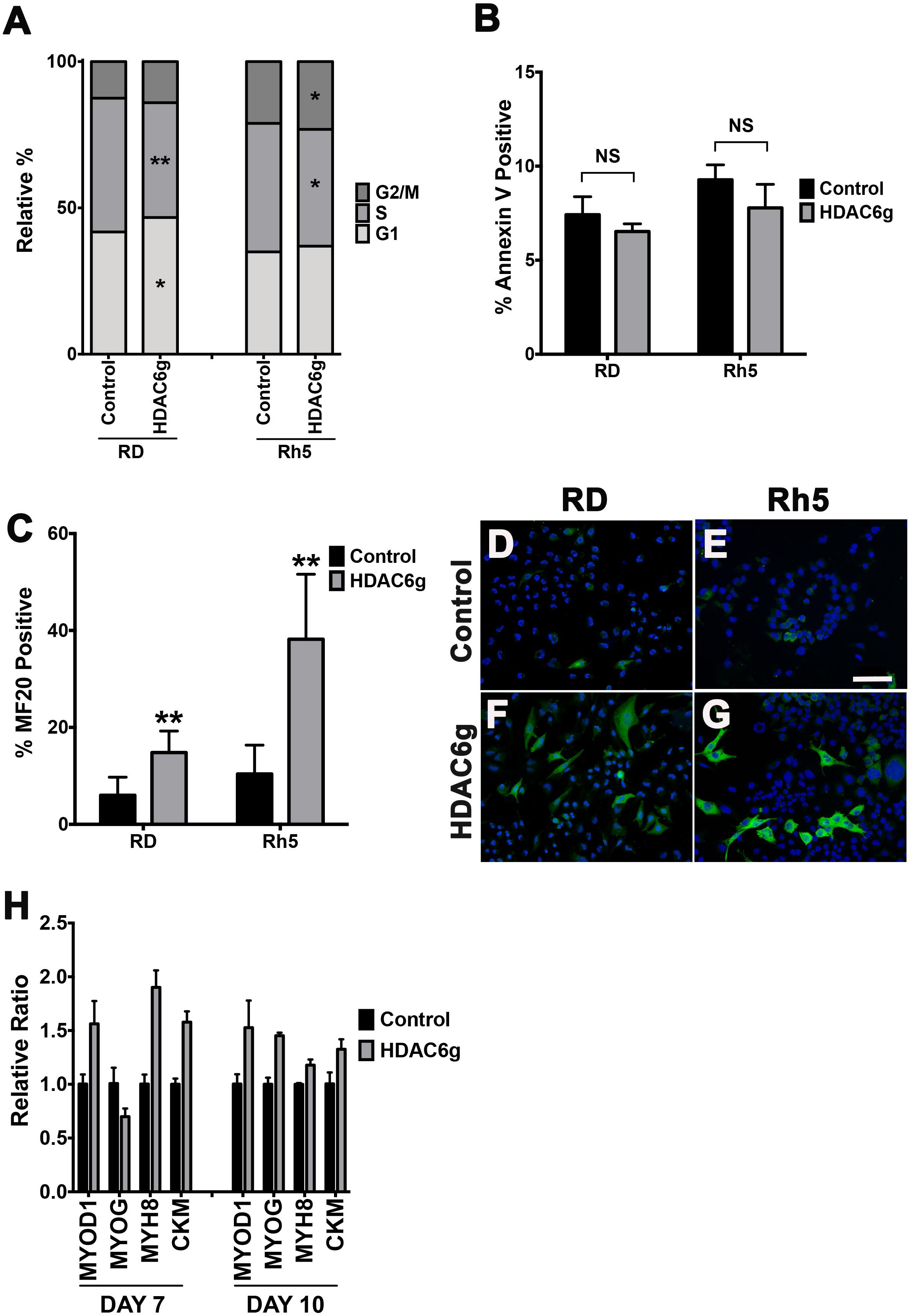
HDAC6 regulates RMS growth by modulating cell cycle progression and tumor cell differentiation. (A) EdU flow cytometry-based cell cycle analysis of RD and Rh5 cells with safe harbor control and CRISPR-mediated *HDAC6* gene disruption. *HDAC6*g = *HDAC6* gRNA. (B) Cell death analysis by Annexin V-based flow cytometry assay in RD and Rh5 cells. NS = No significance statistically by a Student’s t-test. (C) Quantitation of immunofluorescence (IF) against MF20 in RD and Rh5 cells. Summary graphs in A-C are results from 1 representative experiment of 3 independent repeats. Error bar = standard deviation over 3 replicates. * = p < 0.05; ** = p < 0.01. (D-G) Representative IF images in RD and Rh5 cells. Scale bar = 100 microns. (H) Quantitative RT-PCR assessing expression of myogenic genes 7 days and 10 days post tamoxifen-induced CRISPR-mediated *HDAC6* gene disruption in RD cells.

### HDAC6 promotes RMS tumor cell migration and self-renewal

The effects of HDAC6 loss-of-function on the migratory behavior and self-renewal capacity of RMS cells were also assessed. Targeted disruption of *HDAC6* by CRISPR/Cas9 in RD and Rh5 cells resulted in reduced migration by scratch wound healing (Fig. 3 A-B) and transwell assays (Fig. 3 C-D). Using the sphere assay as a surrogate *in vitro* assay for assessing tumor cell stemness and self-renewal^29^, we showed that RD and Rh5 cells with *HDAC6* knockout exhibited reduced frequency and size of sphere formation (Fig. 3 E-F). Expression levels of stem cell markers such as *SOX2* and *NANOG* were also reduced in RD cells with *HDAC6* knockout (Fig. 3 G). Overall, our findings using *in vitro* functional assays suggest that HDAC6 plays an important role in regulating migratory and self-renewal capacity of RMS tumor cells.

**Figure 3.**
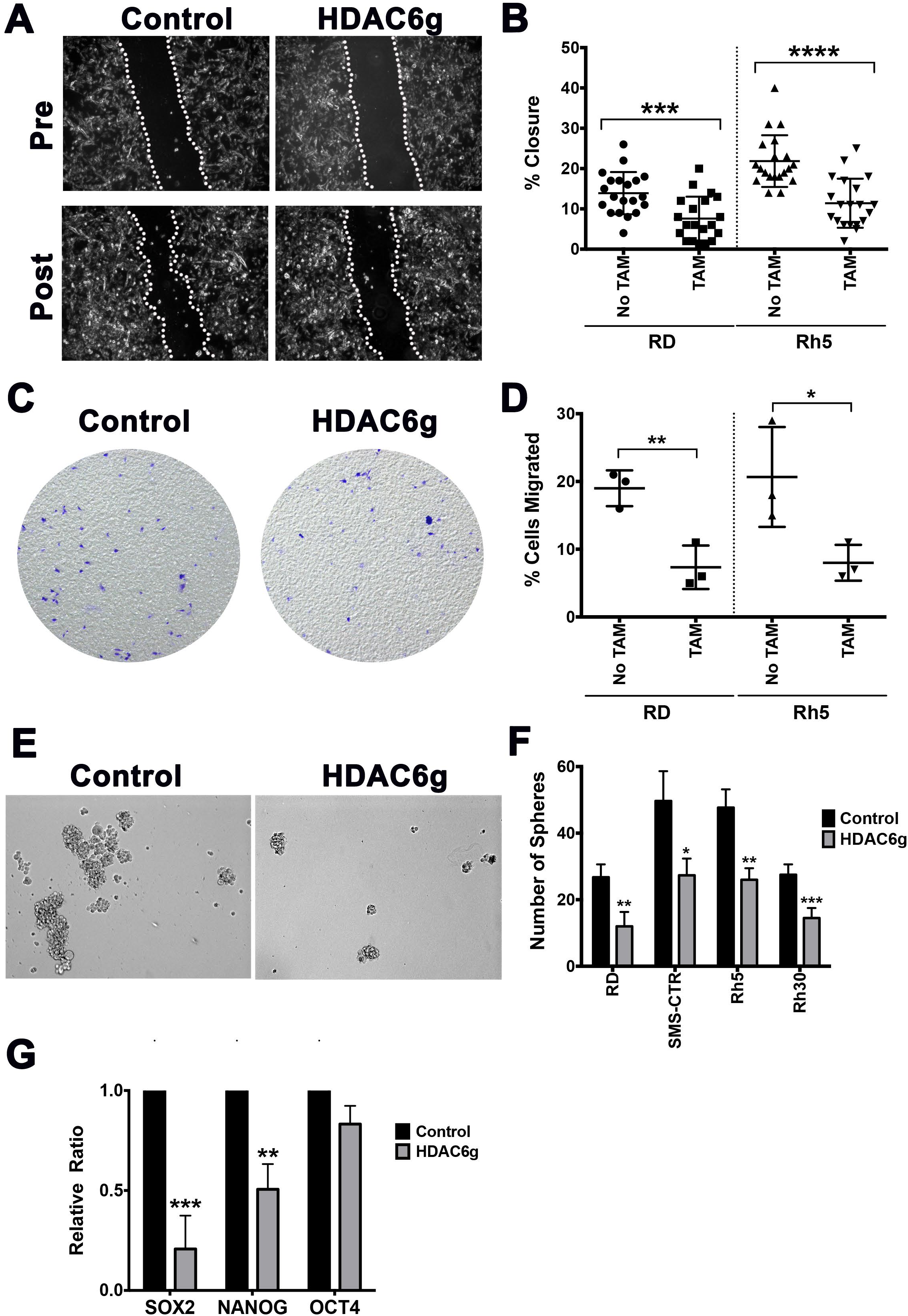
HDAC6 regulates RMS tumor cell migration and self-renewal. (A) Representative images from a wound healing scratch assay in CRISPR safe-harbor control-targeted and *HDAC6* gRNA-targeted RD cells. Post = 18 hours post-scratch. Dashed lines indicate migrating front. (B) Summary of scratch assays in RD and Rh5 with tamoxifen (TAM)-inducible CRISPR *HDAC6* targeting in RD and Rh5. Each datapoint represents a distinct area of the gap. The results shown are from one representative experiment of at least 3 repeats. (C-D) Representative images from a Transwell migration assay. Migrated cells were stained with crystal violet. (D) Summary of Transwell migration assays in RD and Rh5 with tamoxifen (TAM)-inducible CRISPR *HDAC6* targeting in RD and Rh5. Each data point represents a replicate well. Results shown are from one of 3 independent experiments. (E) Representative images from a sphere assay in RD cells. (F) Summary of sphere assays in 4 RMS cells lines. Shown are results of 4 replicates from one of 3 independent experiments. (G) RT-PCR analysis comparing expression levels of stem cell markers in control and *HDAC6*-targeted RD cells. Each error bar in B, D, F and G represents standard deviation. * = p <0.05; ** = p < 0.01; *** = p < 0.001; **** = p < 0.0001.

### Treatment with Tubastatin A recapitulates the HDAC6 loss-of-function effects in RMS

To assess whether treatment of RMS cells with Tubastatin A, a selective HDAC6 inhibitor, could recapitulate the loss-of-function effects of HDAC6 on RMS cells, we first showed that treating a panel of RMS cell lines (RD, SMS-CTR, Rh5 and Rh30) with Tubastatin A *in vitro* significantly inhibited tumor cell growth in a dose-dependent manner (Fig. 4 A-B; Fig. S1 A-B). In scratch wound healing and self-renewal assays, RD and Rh5 cells treated with Tubastatin A (200 nM) showed reduced migration and self-renewal capacity, respectively (Fig. 4 C-F).

**Figure 4.**
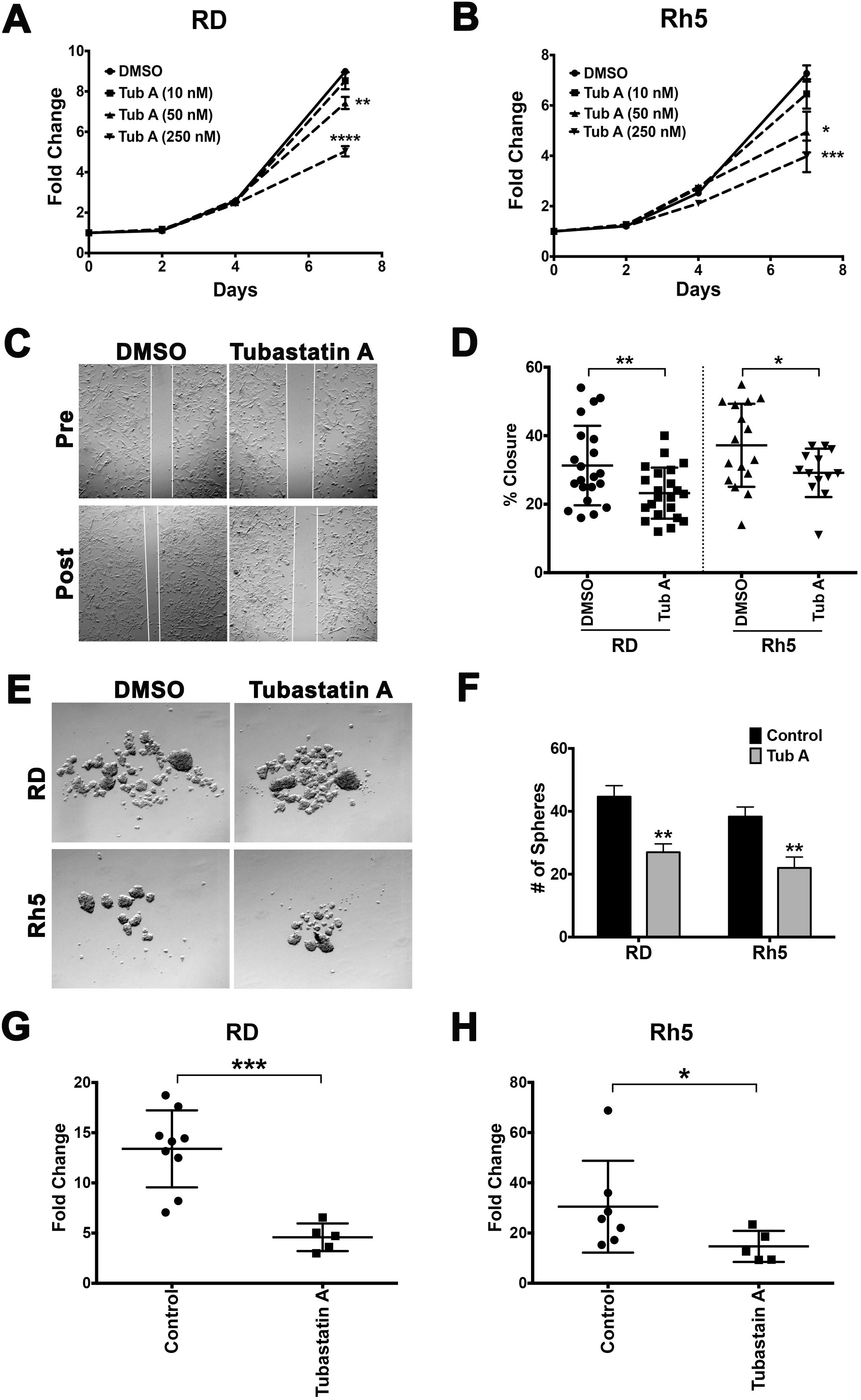
Tubastatin A treatment of RMS cells mimics the HDAC6 loss-of-function phenotypes. (A-B) Cell Titer Glo viability assays assessing the dose-dependent effect of Tubastatin A (Tub A) on cell growth of RD and Rh5 cells over 7 days. Each dose was done in 4 replicate wells. Results from one representative experiment of 3 repeats are shown. (C) Representative images from a scratch assay in RD cells following treatment with DMSO (vehicle) or Tubastatin A (200 nM). (D) Summary of scratch assay analysis in RD cells treated with DMSO or Tubastatin A. Results shown are from one representative experiment of 3 repeats. (E) Representative images of sphere assays in RD and Rh5 cells treated with DMSO or Tubastatin A. (F) Summary of sphere assay analysis in RD and Rh5 cells from one representative experiment of 3 repeats. (G-H) Summary of tumor volume change in xenografts established using RD (G) and Rh5 (H) cells treated with DMSO (vehicle control) and Tubastatin A (10 mg per kg per mouse, drug injection every 3 days over 21 days). Each data point represents a mouse. Each error bar in the graphs of A, B, D, F, G and H represents standard deviation. * = p < 0.05; ** = p < 0.01; *** = p < 0.001; **** = p < 0.0001.

To assess the effect of Tubastatin A treatment in RMS tumor growth *in vivo*, RMS xenografts generated from RD and Rh5 cells were treated with Tubastatin A by intraperitoneal injections at 10 mg per kg per mouse every 3 days for 21 days. RD and Rh5 xenografts treated with Tubastain A showed at least 50% reduction in tumor growth compared to those treated with DMSO as a vehicle control (Fig. 4 G-H; Fig. S1 C-D). RMS xenograft tumors treated with Tubastatin A also showed reduced Ki67 proliferation index based on immunohistochemistry analysis (Fig. S1 E).

To assess whether Tubastatin A treatment affects the self-renewal capacity of RMS cells *in vivo*, we treated zebrafish ERMS tumors with Tubastatin A (10 μM) and DMSO (vehicle) for 7 days and performed limiting dilution transplantation assays. Tumor engraftment was monitored weekly until day 28 post-transplantation. In 3 independent experiments, Tubastatin A-treated ERMS tumors transplanted at limiting dilutions (10^4^, 10^3^ and 10^2^ cells) showed approximately 6-9 fold decrease in self-renewal frequency by Extreme Limiting Dilution (ELDA) analysis^30^ (Table 1). Overall, treatment of RMS cells with Tubastatin A mimics the loss-of-function effects of HDAC6 on tumor cell growth, migration and self-renewal capacity and represents a promising agent for targeted therapy for RMS.

**Table 1.**
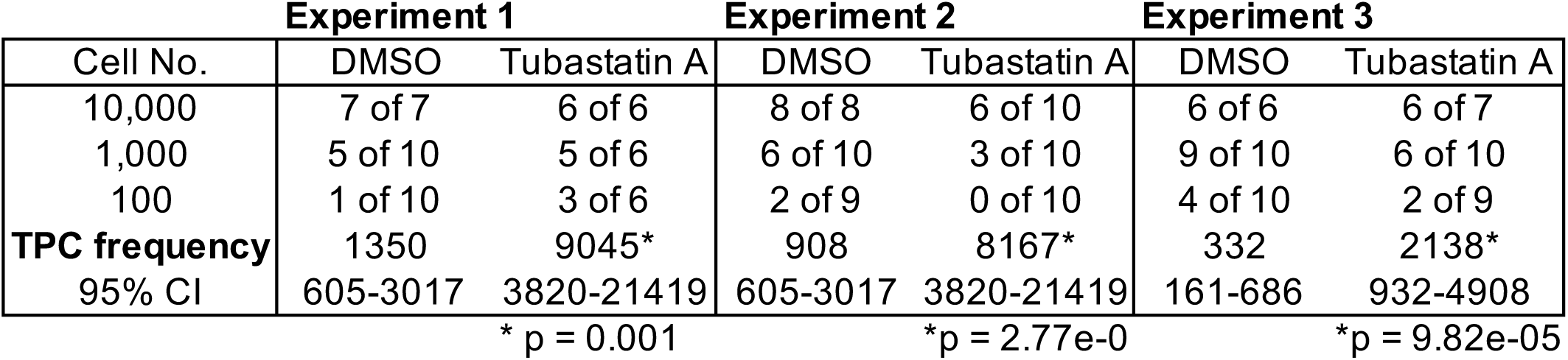
Summary of Limiting Dilution Assays.

### HDAC6 alters the cytoskeletal dynamics to affect RMS cell migration via RAC1

HDAC6 has previously been shown to acetylate the elements of cytoskeleton such as tubulin and actin^20,31^. To assess whether HDAC6 plays a role in modulating the dynamics of cytoskeleton to promote migration of RMS tumor cells, we first showed that there were increased levels of acetylated tubulin in RD and Rh5 RMS cells following targeted disruption of *HDAC6* by CRISPR/Cas9 or treatment with Tubastatin A (Fig. 5 A; Fig. S2 A). Overexpressing either wild-type but not catalytically-dead (cd) Cas9-resistant HDAC6 in RD cells harboring targeted disruption of *HDAC6* alleviated the migratory defect in comparison to overexpression GFP as a control (Fig. 5 B), indicating that the deacetylase activity is required for the HDAC6 function in regulating RMS cell migration. To assess the effects of HDAC6 loss-of-function on actin-dependent cytoskeletal dynamics, we showed by phalloidin staining that there was loss of membrane ruffles, folds and filopodia as well as altered organization of cytoplasmic actin filament in RD and Rh5 cells harboring Cas9-mediated targeted disruption of *HDAC6* following Epidermal Growth Factor (EGF) stimulation in the setting of serum starvation (Fig. 5 C; Fig. S2 B).

**Figure 5.**
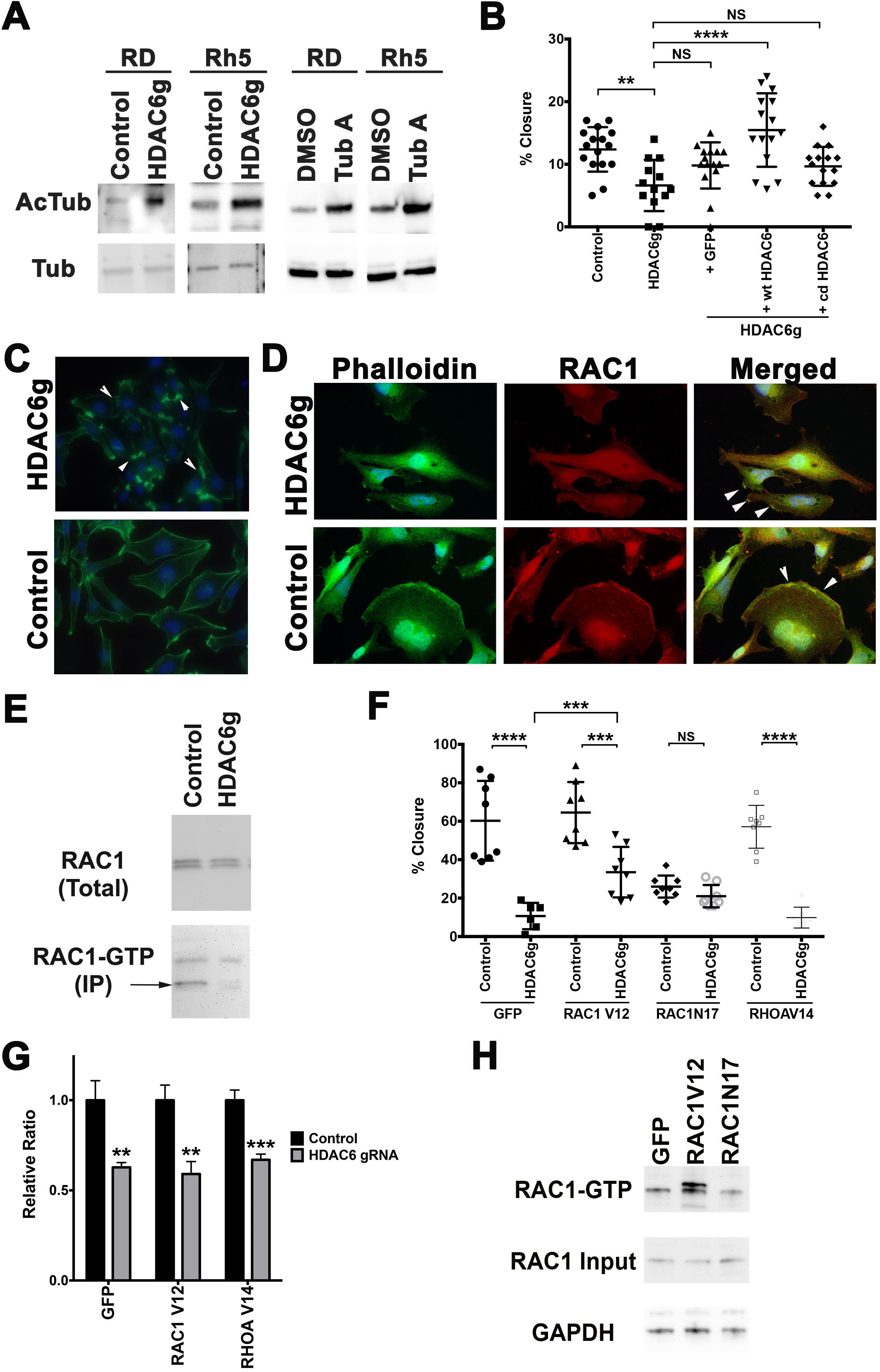
HDAC6 regulates cytoskeletal dynamics to affect RMS cell migration. (A) Western blots against acetylated alpha-Tubulin and alpha-Tubulin in RD and Rh 5 cells with targeted disruption of *HDAC6* by CRISPR or treated with DMSO (vehicle) and Tubastatin A (200 nM). (B) Scratch assay assessing effects of adding back wild-type (wt) HDAC6 and catalytically-dead (cd) HDAC6 in RD cells with *HDAC6* knockout. Results are shown from one representative experiment of at least 3 repeats. (C) Phalloidin staining in RD cells with safe-harbor control region and *HDAC6* CRISPR targeting following serum starvation and 15-minute EGF (50 ng/mL) treatment. Arrowheads point to representative areas of membrane ruffles and filopodia formation. Green = phalloidin, Blue = DAPI. (D) Double IF against HDAC6 (green) and RAC1 (red) in RD cells. (E) RAC1 GTP pulldown assay in RD cells harboring safe-harbor control and *HDAC6* CRISPR targeting. (F) Summary of scratch assays assessing the effects of overexpressing GFP as a control, RAC1V12, RAC1N17 and RHOAV14 in the presence of CRISPR-mediated targeted disruption of *HDAC6* in RD cells following 24 hours of serum starvation and 15 minutes of EGF (50 ng/mL) treatment. (G) Summary of cell growth change by cell counts over 6 days assessing the effects of overexpressing GFP as a control, RAC1V12 and RAC1N17 on cell growth of RD cells harboring targeted disruption of *HDAC6*. Results for the average of 4 replicates for each condition in one of 3 independent experiments are shown. (H) RAC1-GTP pulldown assay in RD cells overexpressing GFP, RAC1V12 and RAC1N17 following 24 hours of serum starvation and 15 minutes of EGF (50 ng/mL) treatment. Each error bar in B, F and G represents standard deviation. NS = no significance, p < 0.05; ** = p < 0.01; *** = p < 0.001; **** = p < 0.0001.

RAC1 is a small GTPase of the Rho family which has been shown to mediate a variety of cellular events including cell adhesion, motility and polarity^32,33^. However, the causal relationship between HDAC6 and RAC1 in mediating the cytoskeletal dynamics to promote cancer cell migration is unclear. To assess whether RAC1 contributes to HDAC6-mediated changes in the cytoskeleton dynamics and cell motility of RMS cells, we first showed that RAC1 and HDAC6 co-localized at the regions of cell membrane ruffles and folds (Fig. 5 D), and that RAC1-GTP levels were reduced in RMS cells with *HDAC6* knockout (Fig. 5 E; Fig. S2 C). Overexpressing the constitutively activated mutant form of RAC1 (RAC1V12) alleviated the migration phenotype (Fig. 5 F) but did not alter the growth phenotype of RD cells with *HDAC6* knockout (Fig. 5 G). A dominant-negative form of RAC1 (RAC1N17) and a constitutively active form of mutant RhoA (RhoAV14), another Rho family GTPase, did not rescue the migration defect in RMS cells with *HDAC6* knockout (Fig. 5 F). Using a RAC1-GTP pull-down assay, RAC1V12 immunoprecipitated increased level of RAC1-GTP compared to GFP control and RAC1N17, supporting the constitutive activity of RAC1V12 (Fig. 5 H). Taken together, HDAC6 alters the microtubule and actin-dependent cytoskeletal dynamics to promote RMS cell migration via RAC1, and there is no cross talk between close members of Rho GTPases in this context.

### RAC1 is essential for RMS cell migration and invasion

As RAC1 is a key mediator of HDAC6 function in regulating RMS cell migration, we next assessed whether RAC1 is essential for RMS cell migration through loss-of-function and gain-of-function studies. Cultured RD and Rh5 RMS cells with targeted disruption of *RAC1* showed reduced migratory capacity by the scratch wound healing assay (p < 0.0001; Figure 6 A-B; Figure S3 A). Similarly, RD and Rh5 cells treated with the selective RAC1 inhibitor, EHop-016, also showed reduced migration (Figure S3 B).

**Figure 6.**
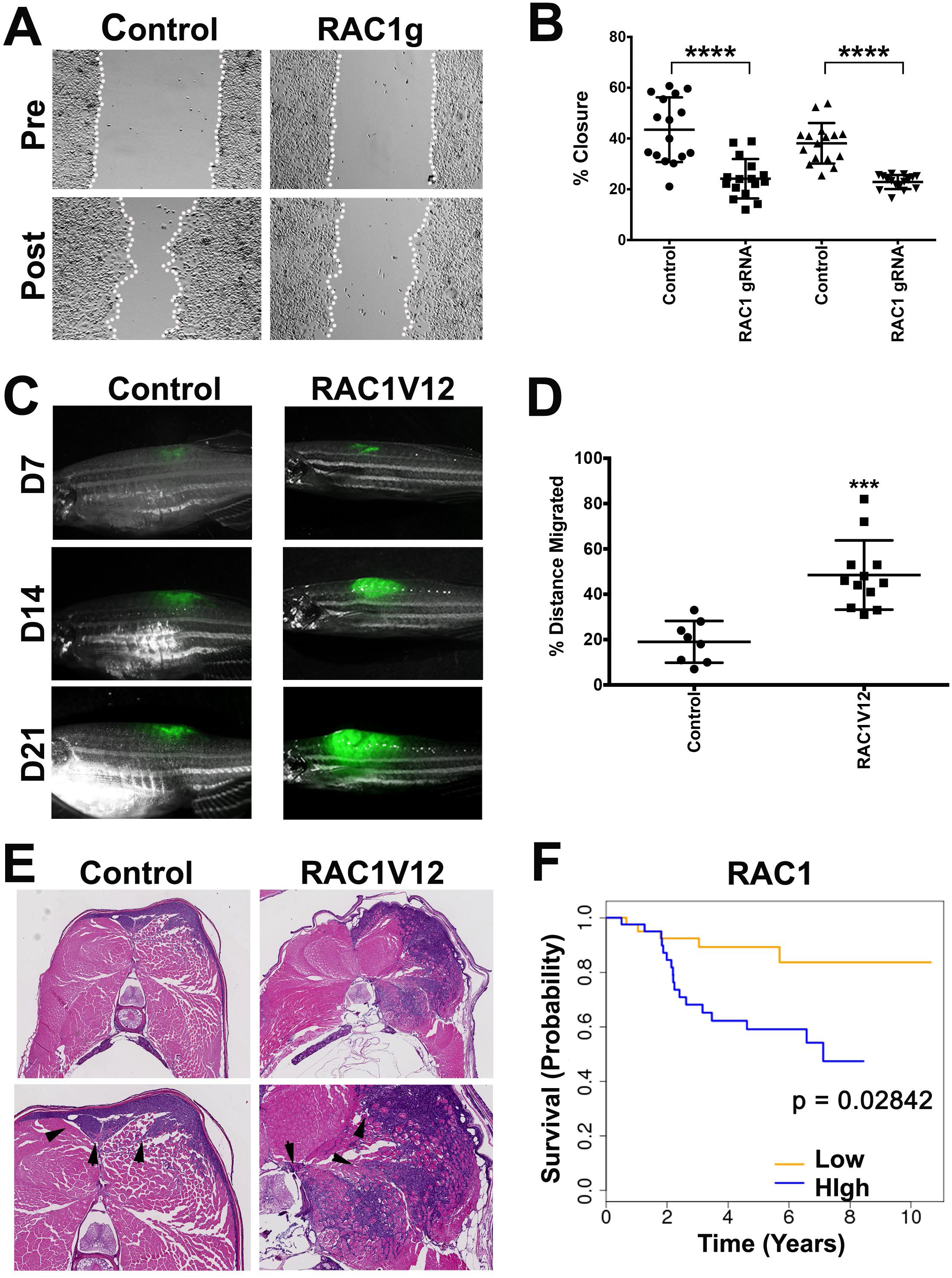
RAC1 is essential for RMS cell migration and invasion. (A) Representative images of RD cells harboring safe-harbor control targeting and *RAC1* gRNA targeting in a wound healing scratch assay. (B) Summary of scratch assays in RD and Rh5 cells with targeted disruption of *RAC1*. The results from one of 3 independent replicate experiments are shown. (C) Representative images of KRASG12D-induced zebrafish ERMS tumors co-expressing GFP and an empty vector (control) or mutant RAC1V12 at day 7, 14 and 21 post-transplantation. (D) Summary of % distance migrated over total dorsal-ventral body axis length for each fish. n = 8 for control and n = 12 for human RAC1V12-expressing tumors. (E) Representative H&E images of control and RAC1V12 expressing zebrafish ERMS tumors. Lower panels show higher magnification with error heads indicating areas of skeletal muscle invasion by tumor cells. (F) Correlation of *RAC1* expression levels with overall survival outcome in RMS patients (63 fusion-negative and 18 fusion-positive). Error bars in graphs of (B) and (D) represent standard deviation. *** = p < 0.001; **** = p < 0.0001.

To assess the gain-of-function effects of RAC1 on RMS cell migration and invasion *in vivo*, KRASG12D-driven zebrafish ERMS tumors co-expressing RAC1V12 or an empty vector as control were transplanted into syngeneic CG1 zebrafish at 10,000 cell per fish via dorsal subcutaneous route, and the fish with engrafted tumors were imaged weekly for tumor cell migration. We showed that by 21 days, zebrafish ERMS tumor cells co-expressing RAC1V12 demonstrate more aggressive growth and increased migration along the dorsal-ventral body axis compared to the control empty vector co-expressing tumor cells (Figure 6 C-D). On histologic analysis, RAC1V12 co-expressing zebrafish ERMS tumors are deeply invasive into the skeletal muscle compared to GFP co-expressing zebrafish ERMS tumors, which showed only superficial invasion in about the same time duration (Figure 6 E).

### RAC1 is associated with poor prognosis in RMS

To assess whether HDAC6 or RAC1 expression is associated with clinical prognosis in RMS, we correlated expression levels of *HDAC6* or *RAC1* mRNA expression in 81 cases with survival data (63 fusion-negative and 18 fusion-positive). Kaplan-Meier curves were generated based on gene expression values dichotomized into over- and under-expressed groups using the median expression value within each cohort as a cutoff. While high expression of *HDAC6* did not correlate with overall survival, high expression of *RAC1* correlated with decreased overall survival (Figure 6 F; log rank test, p = 0.02842, HR = 2.36, 95% CI = 1.095 - 5.084). The findings suggest that expression of *RAC1* can be potentially used as a clinical prognostic indicator in RMS patients.

## DISCUSSION

In this study, we have demonstrated through *in vitro* and *in vivo* functional assays that HDAC6 regulates RMS tumor growth by modulating cell cycle progression and differentiation. HDAC6 also plays an important role in regulating the self-renewal capacity and migration/invasion of RMS cells. RAC1 is an important mediator of HDAC6 function in RMS cell migration/invasion, and high expression of *RAC1* correlates with poor clinical prognosis in RMS patients.

HDAC6 has been shown to deacetylate tubulin, a subunit of microtubule, and cortactin, an actin-associated protein, to affect cytoskeletal dynamics^20,31^. A study in mouse embryonic fibroblasts demonstrates that functional HDAC6 is required for RAC1 activation^23^. However, the functional relationship among HDAC6, the microtubule and the actin filament in driving cancer cell invasion/migration is unclear. In our study, we showed that HDAC6 deacetylase the tubulin subunits of microtubule, and its deacetylase activity is required to promote RMS cell migration. HDAC6 also plays a crucial role in regulating the actin-dependent cytoskeletal dynamics required for RMS cell migration. This is supported by our findings that HDAC6 and RAC1 co-localized in the areas of membrane ruffles and filopodia, which are the changes in the actin elements required for cell motility and invasion, and RMS cells with targeted disruption of *HDAC6* showed reduced formation of membrane ruffling and filopodia. In addition, RAC1 has been shown previously to be required for actin polymerization, stress fiber formation and focal adhesion complex assembly^32^. In this study, loss of HDAC6 resulted in reduced RAC1 GTPase activity, and activated mutant form of RAC1 (RAC1V12) alleviated the migration defect of RMS cells with *HDAC6* knockout. Overall, our findings indicate the deacetylate activity of HDAC6 and activation of RAC1 are essential for the HDAC6-mediated effects on actin-dependent cytoskeletal dynamics required for RMS cell migration and invasion. The crosstalk between the microtubules and actin elements in regulating cellular processes such as cell migration is beginning to be recognized^34^. HDAC6 likely coordinates the direct or indirect interaction between the microtubules and the actin or its associated molecules to promote RMS cell migration and invasion. Further investigation is required to assess whether acetylation of tubulin or other substrates by HDAC6 directly affects RAC1 activation and in turn actin dynamics or HDAC6 independently alters the microtubule and actin dynamics to regulate RMS cell motility.

RAC1 is required for actin cytoskeletal reorganization, a process required for cell migration/invasion during cancer metastasis. Increased expression or activity of RAC1 is associated with metastatic potential in multiple cancer types (e.g. breast, liver and upper urinary tract)^35–37^. However, the functional requirement for RAC1 in cancer cell migration and invasion has not been clearly demonstrated. In this study, we used loss-of-function and gain-of-function studies *in vitro* and *in vivo* to demonstrate that RAC1 is necessary and sufficient for active migration/invasion of RMS cells. High expression of RAC1 also correlates with poor clinical outcomes in RMS patients. Our findings indicate that RAC1 can serve not only as a therapeutic target for reducing invasive and metastatic potential of RMS cells, but also a potential clinical prognostic indicator for predicting RMS patient outcomes.

The self-renewal capacity of tumor initiating cells contributes to cancer relapse and therapy resistance^12^. So far there are limited studies on characterizing the role of HDAC6 in regulating cancer stemness and self-renewal capacity. In glioblastoma, HDAC6 is required for maintaining the glioma stem cell stemness and contributes through its interaction with the Sonic Hedgehog (SHH) signaling pathway to the promote the radio-resistant phenotype^26^. Through *in vitro* sphere assays and *in vivo* limiting dilution assays, we showed that targeted disruption of *HDAC6* by the CRISPR technology and treatment with Tubastatin A, a selective HDAC6 inhibitor, resulted in reduced cancer stemness and self-renewal capacity of RMS cells. RMS spheres generated *in vitro* have been shown to be enriched for a stem cell marker, CD133, and are resistant to treatment with standard-of-care chemotherapy agents^14^. While further investigation is required to determine the pathways modulated by HDAC6 to regulate self-renewal of RMS cells, our findings suggest that targeted therapy using HDAC6 selective inhibitors against cancer stemness represents a promising option for preventing relapse and treatment resistance in RMS.

Our knowledge in the molecular mechanisms underlying RMS self-renewal and metastasis is limited. Using *in vitro* and *in vivo* functional assays, we have characterized the unique role of HDAC6 in regulating RMS tumor growth, self-renewal and migration/invasion. As RAC1 serves as an essential downstream mediator of HDAC6 function in RMS cell migration and invasion, targeting the HDAC6-RAC1 axis will likely improve survival outcomes of RMS patients.

## METHODS

### CRISPR/Cas9 Gene Targeting in human RMS

Single knockout was accomplished by transducing RMS cells with lentiviral virus expressing safe-harbor control or gene-specific double gRNAs and Cas9. Lentiviral transduced RMS cells were plated for cell-based assays 7 days post-transduction following antibiotic selection. Cloning of Cas9 and gRNA expression vectors was performed as described previously^18^.

CRISPR/Cas9 inducible cells were created using piggybac transposition^1^. Stable cell lines were integrated with a construct containing double gRNAs and ERT2-Cas9-ERT2 fusion protein for tamoxifen inducible gene targeting. Inducible gene targeting *in vitro* was accomplished by treating ERMS cells for 3 days with 2 µM 4-hydroxytamoxifen. For structure function studies, the RD cell line harboring the tamoxifen-inducible HDAC6 gRNA/Cas9 cassette were transduced with constructs overexpressing either wild-type or catalytically-dead (cd) HDAC6. The coding portions of wild-type and cd HDAC6 were amplified from the plasmids^38^ obtained from Addgene for cloning purposes. Silent mutations to alter PAM sites to create Cas9-resistant wild-type and cd HDAC6 lentiviral overexpression constructs were introduced using a 4-piece Gibson cloning strategy.

The following gRNAs were used for targeting genes in human RMS cell lines:

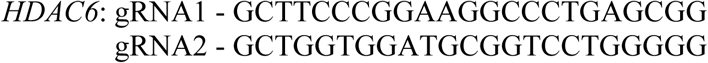

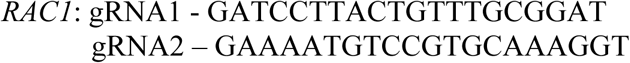

### Assessing Tumor Growth and Self-renewal using Zebrafish ERMS Model

Zebrafish were maintained in a shared facility at the University of Washington under protocol #4330-01 approved by the University of Washington Subcommittee on Research Animal Care. To introduce CRISPR/Cas9-mediated gene targeting in the zebrafish model of KRAS-driven ERMS, a cocktail of DNA constructs containing *rag2*:*KRAS(G12)*-U6-*hdac6* gRNAs, *rag2*:Cas9 and *myog*-H2B:RFP was injected into zebrafish embryos at 1-cell stage. Once RFP+ primary tumors were established (around day 15-20), fluorescent images of tumor fish were taken at tumor onset and 7 days later using a Nikon fluorescent dissecting scope. Tumor volume change was assessed over a 7-day period by measuring tumor area x intensity of fluorescence for each animal using ImageJ as previously described^39^. Deletion of *hdac6* in zebrafish tumors was confirmed by sequencing the PCR products amplified using the primers flanking the gRNA sites.

The *rag2*:*KRAS(G12)*-U6:*hdac6* gRNAs construct was made by using the Gibson cloning strategy to insert the U6:*hdac6* gRNAs cassette into the *rag2*:KRAS(G12) construct obtained from the Langenau lab (Massachusetts General Hospital/Harvard Medical School). The *rag2* promoter and Cas9 coding region were inserted into the *rag2*:Cas9 expression construct using the Gibson cloning strategy.

The following gRNAs were used in zebrafish:

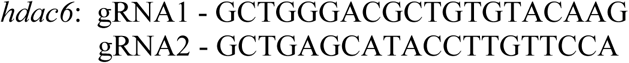

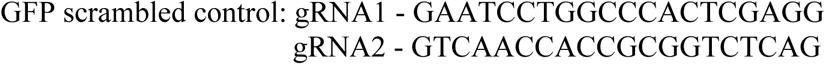

In limiting dilution assays to assess tumor cell self-renewal, zebrafish ERMS tumor cells were transplanted into 3-4 weeks-old juvenile fish and treated with DMSO vehicle or Tubastatin A (10 μM) for 7 days prior to tumor cell harvesting and transplanting in limiting dilutions (10^4^, 10^3^ and 10^2^ cells) into syngenetic adult CG1 zebrafish using the protocol as previously described^39^.

Primary tumors co-expressing KRASG12D, GFP, and empty control vector or RAC1V12 were expanded by transplantation into 3-4 syngeneic adult CG1 fish. GFP-positive tumor cells were isolated by Fluorescent Activated Cell Sorting (FACs) and transplanted subcutaneously into the dorsal region at 10,000 cell per fish. Weekly imaging of tumor-bearing fish post-transplantation was performed using a Nikon fluorescent dissecting scope. The percent distance of the dorsal-ventral body length migrated from the tumor engraftment site over 21 days was quantified using the ImageJ software.

### Cell-based Assays

RMS cell growth was quantified by direct cell counting or the ATP-based CellTiter-Glo luminescent cell viability assay (Promega). To assess myogenic differentiation in RMS, cells were serum starved in 2% horse serum/DMEM for 72 hours prior to fixing in 2% paraformaldehyde for immunofluorescence against MF20 (myosin heavy chain). A flow cytometry-based assay using the Annexin V, Alexa Fluor 647 conjugate (Life Technologies) was used to analyze apoptosis in tumor cells. For cell cycle analysis by flow cytometry, cells pulsed with EdU for 2 hours prior to being harvested and prepped using the Click-iT EdU Alexa Fluor 647 Flow Cytometry Assay kit (Life Technologies). Spheres (“rhabdospheres”) were induced in neurobasal medium enriched with growth factors (EGF, bFGF, PDGF-A and PDGF-B) as previously described^39^. The scratch wound healing assay was performed as previously described^16^.

### Human Xenografts and Drug Treatment

Mouse studies were approved by the University of Washington Subcommittee on Research Animal Care under protocol #4330-01. To establish xenografts in immunocompromised NOD-SCID Il2rg-/- (NSG) mice, approximately 1-2 × 10^6^ RMS cells (Rh5 or RD) were resuspended in Matrigel and injected subcutaneously into the flanks of each 6-7 week-old anesthetized mouse. At tumor onset, tubastatin-A (10 mg/kg) or vehicle (DMSO) was administered by the intraperitoneal route every 3 days for 21 days or until tumor end point. Tumor size was measured by caliper every 3-4 days at tumor onset until tumor end-point or at the end of drug treatment (whichever is earlier). At the end of the experiment, all mice were humanely euthanized for tumor tissue harvesting.

### Immunohistochemistry and Immunofluorescence

A tissue microarray created from archived paraffin tissue blocks for human RMS tumor samples was obtained from Seattle Children’s Hospital. Immunohistochemistry was performed at the Histology and Imaging core facility at University of Washington. The following antibodies including dilutions were used: rabbit polyclonal anti human mouse monoclonal anti human Ki- 67, (1:100, clone: MIB1, Dako), rabbit monoclonal anti human HDAC6 (1:100, clone, D2E5; Cell Signaling), mouse monoclonal anti human RAC1 (1:100, clone 102; BD Biosciences), mouse monoclonal anti acetylated alpha-tubulin (1:200; clone 6-11B-1; Santa Cruz Biotechnology) and mouse monocloncal anti-alpha-tubulin (1:200; B-5-1-2; Santa Cruz Biotechnology). Staining for F-actins was done using Phalloidin conjugated to fluorescent dye 488 (diluted 1:1000; Abnova).

### RAC1 Activation Assay

To assess the levels of RAC1 activation (GTP-bound form) in RMS cells, cultured cells were serum starved in DMEM for 24 hours prior to stimulation with EGF (50 ng/mL) for 10 minutes. Cells were harvested for RAC1-GTP pull down using the Rac1 Activation Assay Biochem Kit (Cytoskeleton, Inc.). Approximately 300 micrograms of protein in each treatment condition was used. Pull down products along with input lysate were run on Western blots to compare the levels of RAC1-GTP to total RAC1 using the antibody against RAC1.

### Western Blots

RMS whole cell lysates prepared in RIPA with protease inhibitors and 2x sample buffer were electrophoresed on a 4-15% gradient SDS-polyacrylamide gel (BioRad) and transferred to PVDF membranes using the TurboTransblot (BioRad). Blots blocked in 5% milk-TBST and probed using the following antibodies and dilutions: HDAC6 (cloneD21B10, Cell Signaling; 1:1000); GAPDH (Cell Signaling; 1:2000); mouse monoclonal anti acetylated alpha-tubulin (1:500; clone 6-11B-1; Santa Cruz Biotechnology), mouse monoclonal anti-alpha-tubulin (1:500; B-5-1-2; Santa Cruz Biotechnology) and RAC1 (1:1000; BD Biosciences). Goat anti-mouse or anti-rabbit HRP conjugated IgG secondary antibodies were obtained from Santa Cruz Biotechnology.

### Survival Association Analysis in RMS patient cohort

The 81 RMS cases with survival and gene expression data were published previously^40^. For survival analysis, p value was calculated based on the log-rank test by R package Survival (https://cran.r-project.org/web/packages/survival/index.html). Note that study subject age and sex, available only on a subset of the data, were not incorporated into the survival analyses because those features are not generally accepted as influencing survival of RMS patients.

### Statistics

Two-tailed Student’s t-test was applied to assess statistical significance in differences between experimental and control samples when appropriate. A p value <0.05 was considered statistically significant.

## ACKNOWLEDGEMENTS

The authors want to thank Terra Vleeshouwer-Neumann and Amy Chen for technical assistance with some experiments. EYC is supported by NIH R01 CA196882. Lin Xu is supported by Children’s Cancer Fund and Rally Foundation.

## AUTHOR CONTRIBUTIONS

EYC contributes to designing and performing the experiments, data analysis and writing the manuscript. TP and KR contribute to performing the experiments and data analysis. LX and SS contribute to data analysis.

