## Supplementary figures and images for "HDAC6 promotes self-renewal and migration/invasion of rhabdomyosarcoma"

### Supplemental Figures

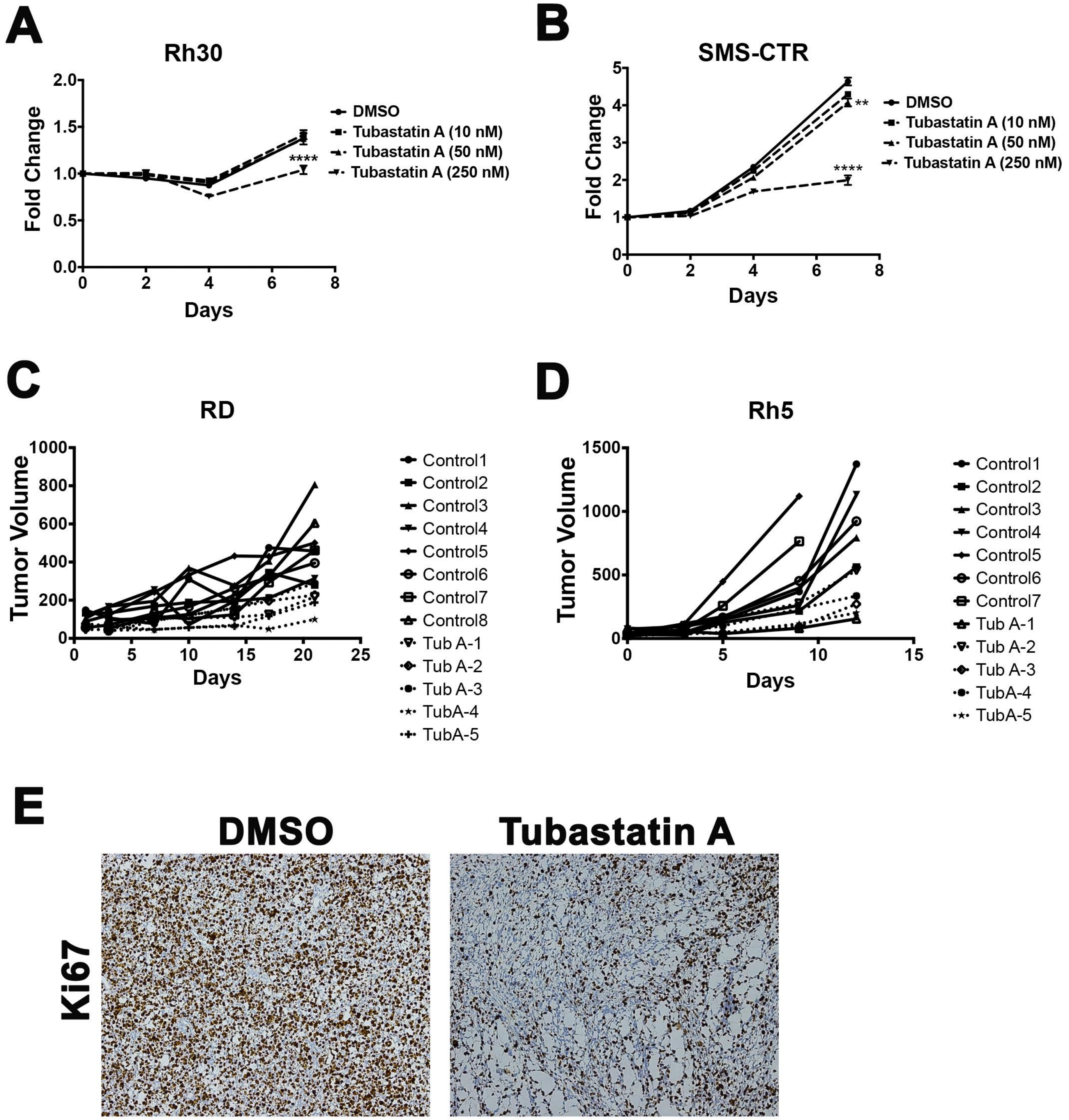

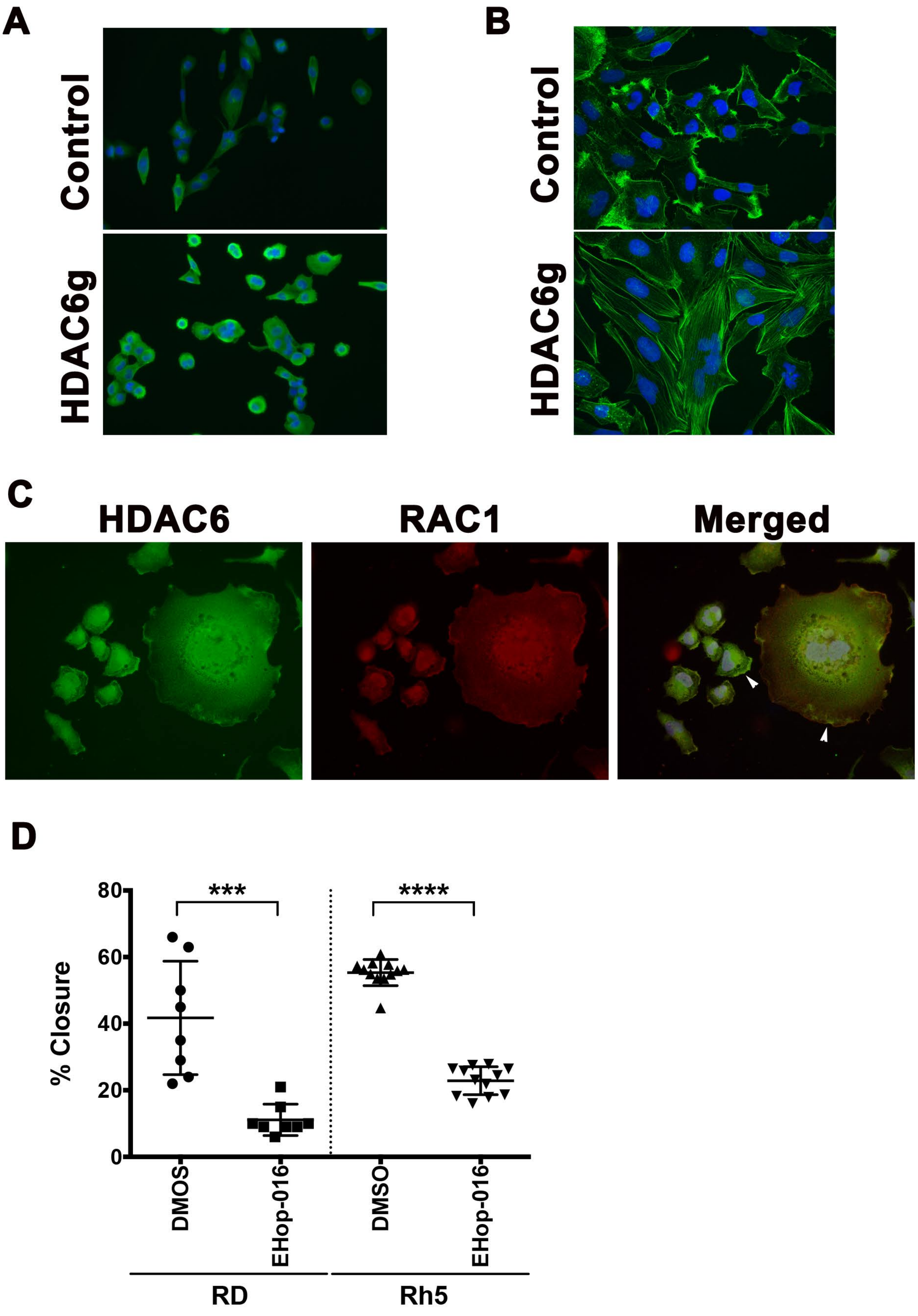

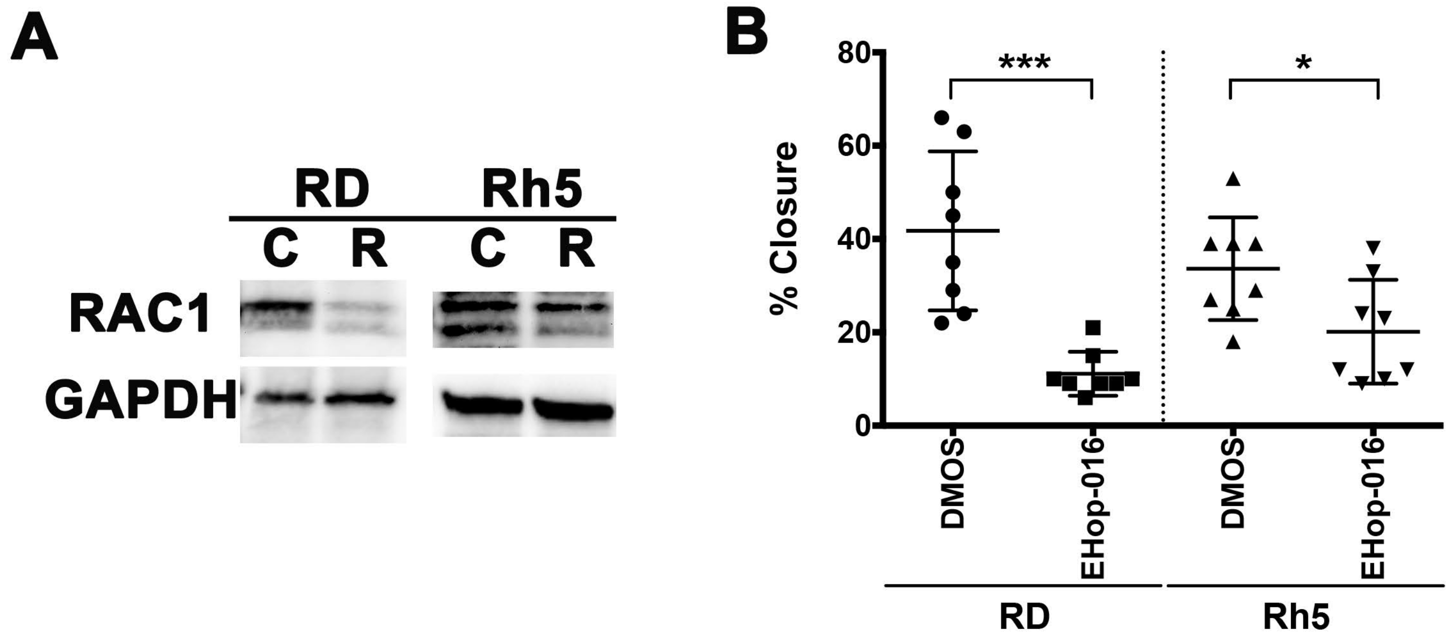
